# Monkey lateral prefrontal cortex subregions differentiate between perceptual exposure to visual stimuli

**DOI:** 10.1101/2024.07.28.605513

**Authors:** Kyoko Leaman, Nadira Yusif Rodriguez, Aarit Ahuja, Debaleena Basu, Theresa H. McKim, Theresa M. Desrochers

**Affiliations:** Department of Neuroscience, Brown University; Department of Biosciences and Bioengineering, IIT Bombay, Mumbai, Maharashtra, India; Department of Biology & Institute for Neuroscience, University of Nevada, Reno; Department of Psychiatry and Human Behavior, Brown University; Robert J. and Nancy D. Carney Institute for Brain Sciences, Brown University

**Author notes:** Correspondence should be addressed to: Theresa Desrochers, 185 Meeting Street, Providence, RI 02912 Box GL-N.

## Abstract

Each day, humans must parse visual stimuli with varying amounts of perceptual experience, ranging from incredibly familiar to entirely new. Even when choosing a novel to buy at a bookstore, one sees covers they have repeatedly experienced intermixed with recently released titles. Visual exposure to stimuli has distinct neural correlates in the lateral prefrontal cortex (LPFC) of nonhuman primates. However, it is currently unknown if this function may be localized to specific subregions within LPFC. Specifically, we aimed to determine whether the posterior fundus of area 46 (p46f), an area that responds to deviations from learned sequences, also responds to less frequently presented stimuli outside of the sequential context. We compare responses in p46f to the adjacent subregion, posterior ventral area 46 (p46v), which we propose may be more likely to show exposure-dependent responses due to its proximity to novelty responsive regions. To test whether p46f or p46v represent perceptual exposure, we performed awake functional magnetic resonance imaging (fMRI) on three male monkeys as they observed visual stimuli that varied in their number of daily presentations. Here we show that p46v, but not p46f, shows preferential activation to stimuli with low perceptual exposure, further localizing exposure-dependent effects in monkey LPFC. These results align with previous research that has found novelty responses in ventral LPFC and are consistent with the proposal that p46f performs a sequence-specific function. Further, they expand on our knowledge of the specific role of LPFC subregions and localize perceptual exposure processing within this broader brain region.

## Introduction

Each day, humans encounter visual stimuli that vary in their perceptual exposure, or how frequently they have previously been seen. Take, for example, browsing the aisles of a bookstore. On each shelf, you are exposed to covers that you have seen many times, which we can consider highly familiar (or having high perceptual exposure). These books may appear alongside recent releases, or books that are more novel (with low perceptual exposure). This visual information may also appear in different contexts, such as a clearly grouped set of books from a popular series versus the books that are scattered across different store displays without evident grouping. Viewing one book after another is one example of what we define as serially presented visual information. When searching a bookstore for a new release, you observe serially presented stimuli while simultaneously maintaining an awareness of the relative familiarity of each title. In this study we will determine the influence of perceptual exposure on neural activity when viewing serially presented stimuli with high or low perceptual exposure.

Perceptual exposure has an effect on neural activity in the lateral prefrontal cortex (LPFC), among other brain regions. Most of the previous work on this topic focuses on novelty and familiarity, which we can consider to be categories on opposing ends of a continuum of different exposure levels. We define novel stimuli as those with limited or no prior perceptual exposure, while familiar stimuli are defined as those that have been experienced extensively (high perceptual exposure). Canonical novelty responses (i.e., higher activation elicited by novel images than familiar ones) have been found across the monkey brain, notably including inferotemporal cortex (IT) (Ghazizadeh et al., 2020; Huang et al., 2018; Ranganath & Rainer, 2003; Zhang et al., 2022), the superior temporal sulcus (STS) (Uhrig et al., 2014), and in LPFC (Ghazizadeh et al., 2020; Ghazizadeh & Hikosaka, 2022; Matsumoto et al., 2007; Mruczek & Sheinberg, 2007; Rolls et al., 2005; Uhrig et al., 2014). These novelty responses are thought to arise, in part, through a graded change in the neural activity, such that novel stimuli have large, distributed neural representations which gradually shift to smaller, more finely tuned sets of neurons as perceptual exposure increases (Koyano et al., 2023; Rainer & Miller, 2000). This process has been specifically recorded in monkey LPFC (Rainer & Miller, 2000). Altogether, it is evident that monkey LPFC responds to differences in perceptual exposure to visual stimuli, showing preferential activation to novel images which decays as exposure increases.

Specific subregions within LPFC may contribute uniquely to these perceptual exposure responses. One such subregion is the posterior fundus of area 46 (p46f) (Rapan et al., 2023), which we have previously identified as tracking visual sequences (Yusif Rodriguez et al., 2023, 2024). A visual sequence consists of a series of images appearing in a given order and can be thought of as one category of serially presented information. We have found that deviations from a known sequence elicit activity in p46f (Yusif Rodriguez et al., 2023, 2024). However, it is unknown whether a similar response may be evoked by infrequently presented stimuli outside of a sequential context. While we currently propose that p46f has a sequence-specific function, this region could respond more broadly to differences in perceptual exposure, as sequence deviants can be thought of as events to which monkeys have a lower exposure. Given that it is unknown to what extent p46f may respond to differences in perceptual exposure outside of the sequential context, we also aim to identify exposure-related effects elsewhere in LPFC as a point of comparison.

Sensitivity to perceptual exposure could differ between even nearby subregions, particularly given the incredible anatomical and functional specificity of LPFC. The posterior ventral subdivision of area 46 (p46v) is an ideal candidate subregion to compare with p46f (Rapan et al., 2023), as it lies adjacent to p46f in the novelty-sensitive ventral LPFC. Previous work has shown that ventral 46 selectively responds to novel images in comparison to familiar images using whole brain fMRI in monkeys (Ghazizadeh et al., 2020) and electrophysiological recordings (Ghazizadeh & Hikosaka, 2022; Matsumoto et al., 2007). Additionally, functional connectivity analyses have identified the specific subregion p46v as being correlated with nearby ventral LPFC region 45a (Rapan et al., 2023), which shows similar novelty responses (Ghazizadeh et al., 2020; Ghazizadeh & Hikosaka, 2022). Thus, we propose that p46v is likely to respond to images with low perceptual exposure, demonstrating a more typical novelty response that may not be shared by p46f.

Here, we investigate whether specific LPFC subregions show activation differences to serially presented stimuli with differing perceptual exposure. We test whether area p46f, which has previously been found to detect deviations in sequential tasks, responds to differences in perceptual exposure. We hypothesize that p46f function is sequence-specific and differences in stimulus exposure alone will not alter its activity. Secondly, we test whether an adjacent subregion, p46v, shows an effect of perceptual exposure, hypothesizing that low perceptual exposure will evoke greater activity than high exposure images, mirroring canonical novelty effects. To address these questions, we used fMRI in awake, behaving monkeys to compare responses to High (∼1500 presentations per day) or Low (∼300 presentations per day) Exposure images. We tested these responses across two timing contexts, one in which images were shown in groups of four (Grouped) and one in which images were presented with jittered time intervals (Ungrouped). We found that p46f did not show preferential activity to low exposure stimuli. We also found that p46v responded more strongly to all stimuli and showed a greater modulation of its response due to perceptual exposure than p46f. Overall, our results demonstrate that p46f does not show activation due to the infrequency of stimuli alone, suggesting that it does not signal relative novelty and is consistent with the possibility that its function is specific to sequences. Additionally, we provide support for p46v as an exposure-sensitive region. These findings expand our understanding of the functional specificity of p46f, an understudied subregion of area 46 that plays a crucial role in daily function, and help to localize perceptual exposure processing within the LPFC at large.

## Methods

### Participants

Subjects for these experiments were 3 adult male rhesus macaques (Monkey W, Monkey J, and Monkey B). Over the course of data collection, they ranged from 6-12 years old and were 9-14 kgs. All procedures followed the NIH Guide for Care and Use of Laboratory Animals and were approved by the Institutional Animal Care and Use Committee (IACUC) at Brown University.

### Task Design and Procedure

For all tasks, the visual stimuli were displayed using an OpenGL-based system created by David Sheinberg at Brown University and controlled by a QNX real-time operating system. In the scanner, stimuli were shown on a 24” BOLDscreen flat-panel display (Cambridge Systems). Eye position was monitored with video eye tracking (Eyelink 1000, SR Research).

For each fMRI scanning session (single day), a new pool of fractals (∼8° visual angle) with randomized coloration and visual features were generated. The new fractals were created using MATLAB scripts from (Kim & Hikosaka, 2013) and instructions from (Miyashita et al., 1991) and luminance matched. Images were presented on a gray background and superimposed with a square fixation spot, which was yellow when the monkey successfully maintained fixation (typically within a 3° window around the square) and red when the monkey failed to fixate.

To encourage fixation, monkeys received juice rewards that were not contingent on the timing of the image presentations and only relied upon maintained fixation. We followed a graduated reward schedule in which sustained fixation resulted in more frequent reward, as outlined by (Leite et al., 2002). Monkeys received the first reward following 4 seconds of sustained fixation. The duration required for juice reward delivery dropped by 0.5 seconds after two rewards of the same fixation length, with the minimum achievable time between rewards being 0.5 seconds. A 0.32 second window was provided for blinks; the monkey’s eyes could leave the fixation point for that duration without triggering a fixation break. Breaking fixation reset the reward schedule to the maximum required fixation to receive reward.

The tasks analyzed here were part of a larger study investigating sequence monitoring that contained four tasks total: a grouped abstract visual sequence task, an ungrouped abstract visual sequence task, non-sequential grouped images, and non-sequential ungrouped images. The grouped sequence task design, reported in (Yusif Rodriguez et al., 2023) and the ungrouped sequence task are not included in the present study. For each scanning session, three new sets of fractals were generated: four used as frequent “habituation” images, three used as infrequent “deviant” images, and four used as “novel” images. Habituation and deviant images appeared in all four tasks in each scanning session, while the novel images only appeared in the non-sequential tasks. In the current experiment, we analyzed activity during presentation of the habituation images (termed High Exposure, presented ∼1500 times per session) and the novel images (termed Low Exposure, presented ∼300 times) in the two non-sequential tasks, Grouped and Ungrouped (**Figure 1B**).

**Figure 1.**
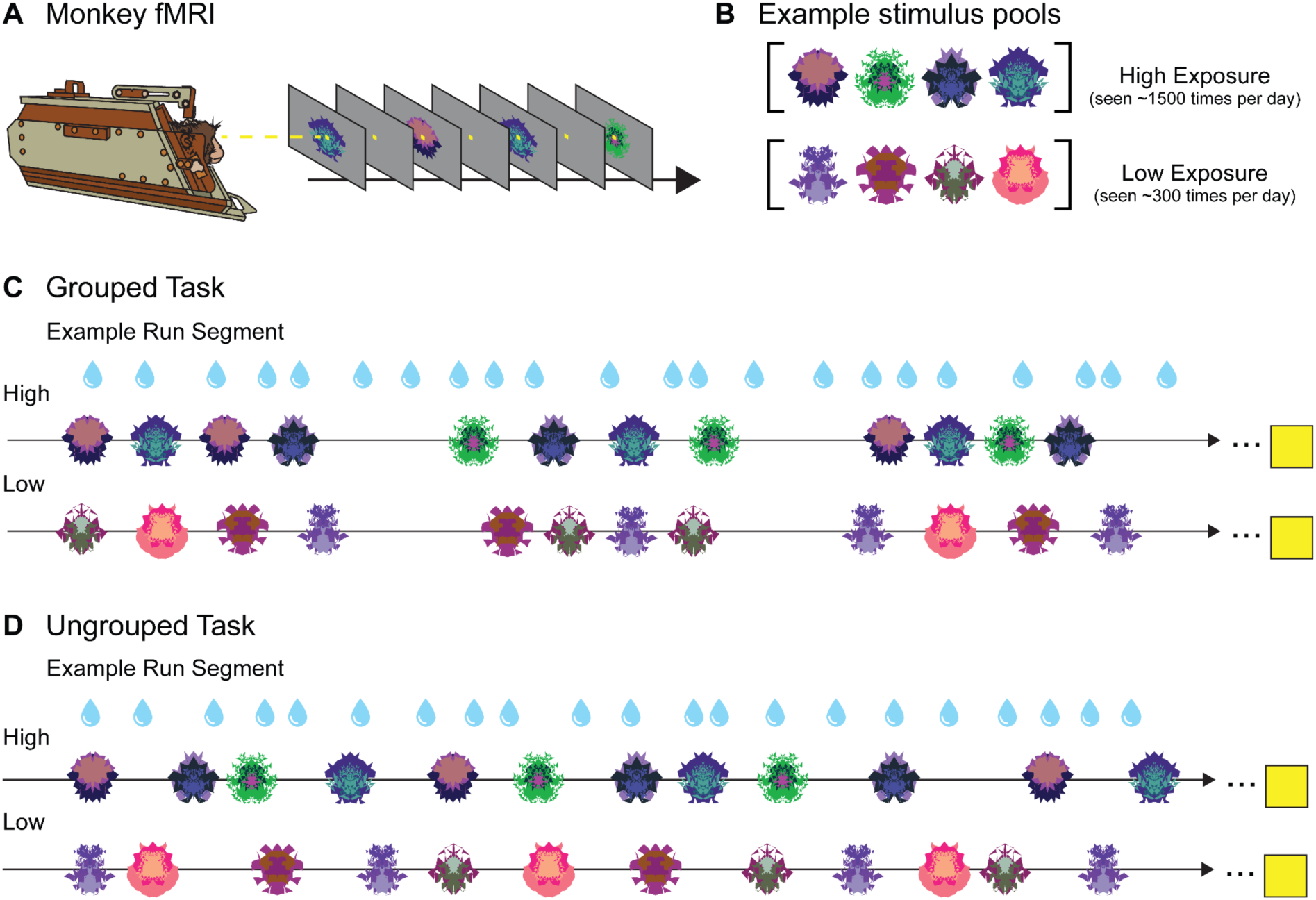
No-report Grouped and Ungrouped viewing tasks. **A.** Monkeys in “sphynx” position fixate on central fixation square during fMRI scanning for both tasks. **B.** Example High and Low Exposure stimulus pools for both Ungrouped and Grouped tasks depict images that would be used for a single scanning session. Over the course of a standard scanning session, High Exposure images will be presented ∼1500 times on average while Low Exposure images will be presented ∼300 times on average. **C.** Example partial Grouped task run showing two of three possible block types (snippets of High and Low Exposure blocks are shown, the third deviant block type is not reported here). Three pseudorandomized four-item example groupings are shown for a High Exposure block (top row) and Low Exposure block (bottom row). Four-item image groups appear in six possible timing templates. All blocks contained 30 four-item groupings (120 total image presentations). The yellow square indicates the 14 second fixation period between each block. **D.** Example partial Ungrouped task run showing snippets of High and Low block types (the third is not shown). Twelve pseudorandomized image presentations with jittered timing are shown for a High Exposure block (top row) and Low Exposure block (bottom row). All blocks contained 120 image presentations. The yellow square indicates the 14 second fixation period between each block. Blue water droplets schematize reward delivery, which is decoupled from image events and delivered on a graduated schedule based on the duration the monkey has maintained fixation.

#### Grouped Task

Images in the Grouped Task were presented in groups of four displayed in a pseudorandom order that ensured no consecutively repeated images. In the present study, there were six possible timing templates and two stimulus pools (High and Low Exposure, **Figure 1C**). Deviant timing templates and image types were present in the task but are not described here because those blocks were not included for analysis.

Total duration for a four-item group was either short (1.1 s), medium (1.7 s), or long (2.3 s). Within each timing template, there were two possible image durations: 0.1 s and 0.2 s for short, 0.1 s and 0.3 s for medium, and 0.2 s or 0.3 s for long. Inter-stimulus intervals ranged from 0.1 s to 0.5 s based on the timing that would evenly space the stimuli within the total sequence duration. Intervals between groups were jittered, with a mean of 2 seconds and a range of 0.25 - 8 seconds.

All blocks contained 30 four-item groups (120 total images). There were four block types, only two of which were analyzed here. The first block contained images drawn from the High Exposure pool and each four-item group used one of the six timing templates, such that each timing scheme appeared five times in total. In the second block analyzed, all timing templates remain the same but images were from the Low Exposure pool. Each run contained four blocks interleaved with 14 s fixation periods. The two blocks presented in this study were the first and the fourth of these blocks; the middle blocks are presented elsewhere (Yusif Rodriguez et al., 2024). Runs lasted approximately 10 minutes and monkeys typically finished 2-4 runs of this task (along with other tasks) in one scanning session. A run was initiated when the monkey successfully fixated for ∼4 juice rewards (12-16 s) during the pre-scan period.

#### Ungrouped Task

Images in the Ungrouped Task were presented in a pseudorandom order and without coherent grouping in time (**Figure 1D**). All stimuli had a 0.2 second duration, with a variable inter-stimulus interval that was jittered between 0.25 and 8 seconds (mean 2 s). The same stimulus pools (High and Low Exposure) were used, along with the deviant images that are not reported in the present study.

There were 120 images per block presented pseudorandomly such that the same fractal could not appear twice consecutively. One run contained 3 blocks appearing in a specified order, with 14 s of fixation between each. The first contained only High Exposure images, the second contained 80% High Exposure images and 20% deviant images (not analyzed here), and the third contained only Low Exposure images. The average duration of an entire block was approximately 120 seconds. A single run took roughly 15 minutes to complete, and monkeys completed about 2-4 runs of this task (along with others) per scanning session. As with the Grouped Task, a run was initiated by the monkey successfully fixating.

### fMRI Data Acquisition

Details of data acquisition for these experiments, including sample size justification analyses, have previously been described in (Yusif Rodriguez et al., 2023) and will be briefly outlined here.

Monkeys were positioned in a custom MR-safe primate chair (Applied Prototype, Franklin, MA or built at Brown University) with their head restrained by a plastic headpost (PEEK, Applied Prototype, Franklin, MA). They were trained in the “sphynx” position facing the screen. Monkeys were habituated to the injection of contrast agent, recorded MRI sounds, being transported to the MRI facility, and wearing earplugs (Mack’s Soft Moldable Silicone Putty Earplugs, Kid Size) prior to the start of scanning sessions. Monkeys trained on the tasks using a different set of stimuli than those generated at the scanner.

A contrast agent is used to increase the contrast-to-noise ratio of acquired fMRI data (Leite et al., 2002). Monkeys received an intravenous contrast agent injection 30 minutes to an hour prior to each scanning session. The contrast agent used was monocrystalline iron oxide nanoparticle (MION, 30 mg per mL Feraheme (ferumoxytol), AMAG Pharmaceuticals, Inc., Waltham, MA or 30 mg per mL BioPal Molday ION, Biophysics Assay Lab Inc., Worcester, MA). Seven mg/kg of MION was injected into the saphenous vein below the monkey’s knee and then flushed with double that volume of sterile saline. No additional MION was administered during scanning.

All scans were performed at the Brown University MRI Research Facility using a Siemens 3T PRISMA MRI system and a custom-built six-channel surface coil (ScanMed, Omaha, NE). Consistent with training, awake monkeys performed no-report viewing tasks in the scanner while sitting in “sphynx” position and with their heads restrained (**Figure 1A**). Functional scans were conducted using a fat-saturated gradient-echoplanar sequence with a repetition time (TR) of 1.8 s, an echo time (TE) of 15 ms, 80° flip angle, 40 interleaved axial slides, and 1.1 x 1.1 x 1.1 mm voxels. Anatomical scans were also collected: a T1-MPRAGE (TR 2700 ms, TE 3.16 ms, flip angle 9°, 208 sagittal slices, 0.5 x 0.5 x 0.5 mm voxels), a T2 anatomical (TR 3200 ms, TE 410 ms, variable flip angle, 192 interleaved transversal slices, 0.4 x 0.4 x 0.4 mm voxels), and an additional high resolution T2 anatomical (TR 8020 ms, TE 44 ms, flip angle 122°, 30 interleaved transversal slices, 0.4 x 0.4 x 1.22 mm voxel).

### fMRI Data Analysis

The data analysis and statistical modeling parameters used here are largely the same as those described in (Yusif Rodriguez et al., 2024). All analyses were performed in MATLAB (Mathworks, R2017b) and used SPM 12 (http://www.fil.ion.ucl.ac.uk/spm). Preprocessing steps were completed run-by-run and included reorientation in the x, y, and z dimensions, realignment/motion correction, coregistration/normalization (normalized to the 112RM-SL macaque atlas from (McLaren et al., 2009), and spatial smoothing (using a 2mm isotropic Gaussian kernel, run on gray and white matter separately). The T1-MPRAGE was skull stripped using the FSL BET brain extraction tool (http://www.fmrib.ox.ac.uk/fsl/) prior to normalization. For a run to be included in analysis, both the fixation data and the acquired volumes had to pass a data quality check. A run was excluded if the monkey fixated for less than 80% of the time. The data quality of the volumes was checked using the ART Toolbox (Artifact Detection Tools, https://www.nitrc.org/projects/artifact_detect). A volume was excluded from further analyses if there was greater than 1 voxel (1.1 mm) motion in any direction and runs that had more than 12% of their volumes excluded were removed from analysis (**Table 1**).

**Table 1.** Data included and excluded from analyses.

| Percent Excluded Fixation |  |  |  |  |  |
| --- | --- | --- | --- | --- | --- |
|  | Monkey B |  | Monkey J | Monkey W |  |
| Ungrouped | 7.78% |  | 10.6% | 4.44% |  |
| Grouped | 6.96% |  | 10.13% | 5.06% |  |
| Percent Excluded Motion |  |  |  |  |  |
|  | Monkey B |  | Monkey J | Monkey W |  |
| Ungrouped | 13.3% |  | 1.11% | 1.67% |  |
| Grouped | 16.5% |  | 0.63% | 1.89% |  |
| Total Included Runs |  |  |  |  |  |
|  | Monkey B | Monkey J | Monkey W |  | Total Runs |
| Ungrouped | 27 | 35 | 55 |  | 117 |
| Grouped | 17 | 38 | 43 |  | 98 |

Separate models were constructed for the Grouped and Ungrouped tasks. For both models, data were binned to distribute runs from different scanning sessions into pseudo-subject bins that each contained data from a single monkey. For each task, 10 bins were created, each containing ∼10-12 runs. Bins were pseudo-randomly constructed to balance the distribution of runs from early and late scanning sessions. Within-subject statistical models were run in SPM 12 using the general linear model (GLM) assumptions for each pseudo-subject bin. Condition regressors were convolved with a gamma function which modeled MION hemodynamics (shape parameter of 1.55 and scale parameter of 0.022727).

Nuisance regressors in both models included motion, reward, image variability (ART Toolbox, standard deviation of motion variability), and outlier volumes (ART Toolbox, scan-to-scan motion and global signal change used for outlier detection; global signal outlier threshold of 4.5 and a motion threshold of 1.1mm). A regressor was included with a “1” placed at the position of each outlier volume. The first 24 seconds of a run and reward times were also included as nuisance conditions. Regressors were estimated with a fixed effects model applied to each bin. Bin-specific effects in the whole brain estimate were then entered into second-level analyses with bin included as a random effect. We determined significance with one-sample t-tests (contrast value vs zero) with a significance threshold of 0.005. Multiple comparison corrections were performed using the false discovery rate (FDR) error correction (p < 0.05).

In the Grouped task, regressors of interest were constructed as previously described in (Yusif Rodriguez et al., 2024) (No Sequence task) and briefly described here. The first image in each group of four images was modeled as an instantaneous onset. High and Low Exposure image groups were modeled separately for the short, medium, and long timing templates, yielding six primary regressors of interest. High and Low Exposure images were in blocks one and four of the Grouped task. Blocks two and three were also modeled as follows, but not included in analyses. Instantaneous onsets for each group of images were again modeled separately for each timing template which included frequent short, medium, and long groupings as in the other blocks, and infrequent “deviant” timing templates (medium with 0.2s stimulus duration, two-item, and six-item groupings). Single images that were used in deviant sequences in other tasks were mixed in with the groupings (20% of total images presented) and a separate onset regressor was included for those images. Thus, seven total regressors were included to model the blocks not included for analyses. Each run of the Grouped task contained 13 total regressors.

In the Ungrouped task, instantaneous onsets were modeled for each individual image. High and Low Exposure images were modeled separately. Images previously used as deviants in separate sequence tasks that occurred 20% of the time in the middle block were also modeled as separate instantaneous onsets, but not included for subsequent analyses. There were three total regressors in the Ungrouped task.

The area 46 subregion ROIs were acquired from the MEBRAINS Multilevel Macaque Atlas (Balan et al., 2024). The subregion image warps were shifted from their native space to 112RM-SL space with Rhemap (Sirmpilatze & Klink, 2020) (https://github.com/PRIME-RE/RheMAP). Individual warps were applied to create the images used for the p46v ROI. To best match the sequence-monitoring ROI used in (Yusif Rodriguez et al., 2023), which spanned p46df and p46vf, these subregions were combined to generate the p46f ROI (p46df + p46vf). We specifically examined responses in the right hemisphere, as sequence responses were previously observed in the right hemisphere specifically (Yusif Rodriguez et al., 2023).

To compare the level of activation in response to stimulus exposure within and across ROIs while controlling for variance, the T-values for each pseudo-subject bin were calculated from the condition of interest > baseline contrasts using the Marsbar toolbox in SPM 12 (Jean-Baptiste Poline, 2002). These T-values were entered into two-way repeated-measures ANOVAs with monkey identity included as a covariate.

## Results

Three male monkeys (*Macaca mulatta*) completed no-report image viewing tasks while awake in an fMRI scanner (**Figure 1**). Presented fractal images were either High Exposure, meaning that the monkeys were exposed to them numerous times in the context of multiple tasks over the course of a scanning session (more than 1500 exposures to each image on average), or Low Exposure, meaning they were presented less often and across fewer tasks (around 300 exposures to each image on average). Responses to High and Low Exposure images were analyzed in two task contexts: Grouped and Ungrouped. During the Grouped task, fractals were shown in groups of four and appeared in a structured timing template, while in the Ungrouped task the image timing was jittered. We compared neural responses to High and Low Exposure images to investigate the effect of perceptual exposure on activity in monkey area 46 subregions in both Grouped and Ungrouped timing schemes. All statistical tests were performed on pseudo-subject bins (n = 10 bins for Grouped and 10 bins for Ungrouped, ∼10-12 runs per bin). For each condition, t-values for condition > baseline in each ROI were extracted and entered into statistical tests along with a covariate for monkey identity (n = 3). As we do not have questions regarding individual differences, our discussion focuses on the condition effects, although we report between-monkey variation.

### The fundus of area 46 does not preferentially respond to Low Exposure stimuli

Our first aim was to determine whether perceptual exposure modulates responses in p46f. This region was previously identified as responding to deviations from a sequence (Yusif Rodriguez et al., 2023, 2024), but the extent of its role in processing perceptual exposure outside of a sequence was unknown. To address this question, we compared right p46f responses to High Exposure images (seen ∼1500 times in a day) and Low Exposure images (seen ∼300 times in a day) in the Grouped and Ungrouped tasks (**Figure 1**). There was a marginal difference between High Exposure and Low Exposure image responses in p46f, but perhaps contrary to typical low exposure (novelty) responses the activation was numerically greater for High Exposure than Low Exposure (p = 0.09, **Table 2**, **Figure 2**).This subregion showed no differences between the Grouped and Ungrouped tasks (p = 0.92) and no interaction between task and image exposure (p = 0.9). These results suggest that p46f does not show the proposed canonical novelty response to Low Exposure stimuli but may rather show preferential activation for more frequently seen images, although this effect is marginal.

**Figure 2.**
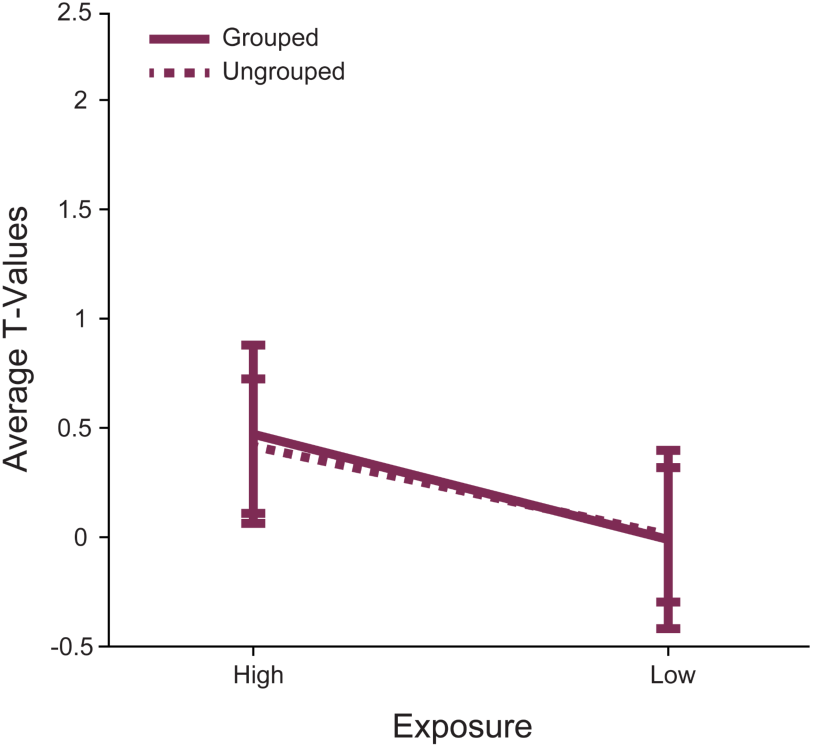
Right area p46f does not respond differently to Low Exposure versus High Exposure images. The mean T-values for each condition > baseline are shown. The error bars represent the 95% confidence intervals (1.96 x standard error of the mean). Responses to stimuli in the Grouped task are shown in a solid line, while responses in the Ungrouped task are shown with a dashed line. Responses to High and Low Exposure stimuli were different such that there was a marginal main effect of stimulus exposure.

**Table 2.** Comparison of activity in right p46f for exposure levels across both tasks using repeated measures ANOVA. Bolded p-values are conditions of interest.

| Factor | DFs | F | p | $\eta^2$ |
| --- | --- | --- | --- | --- |
| monkey | 2,16 | 1 | 0.38 | 0.12 |
| grouping | 1,16 | 0 | 0.92 | 0 |
| exposure | 1,16 | 3.2 | **0.09** | 0.17 |
| monkey:exposure | 2,16 | 0.3 | 0.76 | 0.03 |
| grouping:exposure | 1,16 | 0 | 0.9 | 0 |

### Areas p46v and p46f show distinct response patterns due to exposure level and grouping

Next, we investigated whether a neighboring area 46 subregion responded distinctly to image exposure. Prior work has identified ventral 46 as showing novelty responses (Ghazizadeh et al., 2020; Ghazizadeh & Hikosaka, 2022; Matsumoto et al., 2007). This led us to select p46v, which lies adjacent to p46f, for comparison. We predicted that responses would differ between p46f and p46v, expecting to see preferential p46v activation to Low Exposure stimuli. We directly compared p46v and p46f responses to High and Low Exposure images in Grouped and Ungrouped timing contexts. We found that responses were overall greater in p46v than in p46f in the Grouped context (p < 0.001, **Table 3**, **Figure 3A**). Additionally, p46v showed greater responses to Low than High Exposure images, resulting in an interaction when comparing areas (p = 0.03). In the Ungrouped context, p46v once again showed significantly greater responses than p46f (p = 0.01, **Table 4**,**Figure 3B**), and there was a marginal main effect of perceptual exposure (p = 0.09). Unlike the Grouped task, there was no interaction between exposure and ROI in the Ungrouped task, which could indicate that responses in p46v differ due to image grouping. To test this possibility, we compared p46v responses in the Grouped and Ungrouped tasks. We found a marginal interaction, such that the difference between Low and High Exposure items is marginally greater in the Grouped compared to the Ungrouped task (p = 0.12, **Table 5**). These results indicate that responses in p46f and p46v are different, with p46v showing a tendency to respond more to Low Exposure items in the Grouped task.

**Figure 3.**
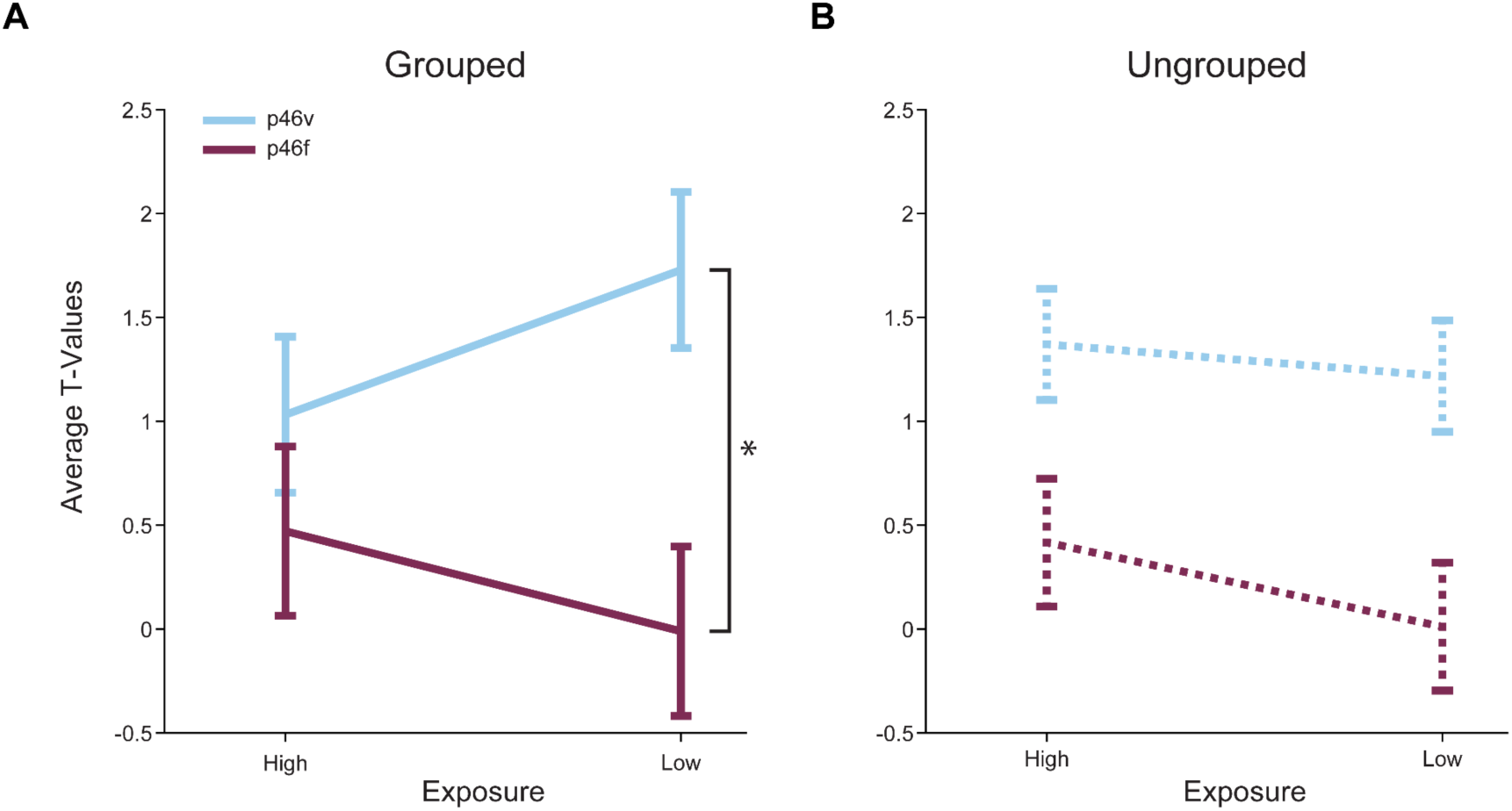
Right areas p46v and p46f respond differently across conditions in both tasks. The mean T-values for each condition > baseline are shown. The error bars represent the 95% confidence intervals (1.96 x standard error of the within-bin mean). Responses in p46v are shown in blue and p46f is shown in purple. **A.** High Exposure compared to Low Exposure images in the Grouped task show a significant interaction effect with ROI (indicated by asterisk, p = 0.05) and a highly significant main effect of ROI. **B.** High Exposure compared to Low Exposure images in the Ungrouped task show a significant main effect of ROI and a marginal main effect of exposure.

**Table 3.** Comparison of activity in the Grouped task for exposure levels across both ROIs using repeated measures ANOVA. Bolded p-values are conditions of interest.

| Factor | DFs | F | p | $\eta^2$ |
| --- | --- | --- | --- | --- |
| monkey | 2,16 | 16.9 | 0 | 0.68 |
| ROI | 1,16 | 47.2 | **0** | 0.75 |
| exposure | 1,16 | 0 | 0.96 | 0 |
| monkey:exposure | 2,16 | 4.3 | 0.03 | 0.35 |
| ROI:exposure | 1,16 | 6 | **0.03** | 0.27 |

**Table 4.** Comparison of activity in the Ungrouped task for exposure levels across both ROIs using repeated measures ANOVA. Bolded p-values are conditions of interest.

| **Factor** | **DFs** | **F** | **p** | **$\eta^2$** |
| --- | --- | --- | --- | --- |
| monkey | 2,16 | 3.5 | 0.05 | 0.31 |
| ROI | 1,16 | 7.6 | **0.01** | 0.32 |
| exposure | 1,16 | 3.2 | **0.09** | 0.17 |
| monkey:exposure | 2,16 | 2.9 | 0.09 | 0.26 |
| ROI:exposure | 1,16 | 0.4 | 0.51 | 0.03 |

**Table 5.** Comparison of activity in right p46v for exposure levels across both tasks using repeated measures ANOVA. Bolded p-values are conditions of interest.

| **Factor** | **DFs** | **F** | **p** | **$\eta^2$** |
| --- | --- | --- | --- | --- |
| monkey | 2,16 | 20.8 | 0 | 0.72 |
| grouping | 1,16 | 0.1 | 0.72 | 0.01 |
| exposure | 1,16 | 0.9 | 0.36 | 0.05 |
| monkey:exposure | 2,16 | 0.6 | 0.57 | 0.07 |
| grouping:exposure | 1,16 | 2.7 | **0.12** | 0.15 |

### Low Exposure images evoke activity in canonical novelty-sensitive brain regions

Whole brain analyses supported the ROI results and showed Low Exposure responses in known novelty-sensitive regions. Though the number of times the monkey views the Low Exposure stimuli is relatively few compared to High Exposure images, the images are only truly novel (in that they have never been seen before) the first time they are viewed in a session. To determine whether our Low Exposure images might be comparable to novel stimuli used in other studies, we conducted whole-brain voxel-wise contrasts comparing Low and High Exposure items for both the Grouped and Ungrouped tasks. In the Grouped task Low Exposure > High Exposure, there were significantly greater responses in regions TEO, TEpd, 45/46v, and 13l (FDRc < 0.05, height p < 0.005 unc., extent = 116; **Table 6**, **Figure 4**). All of these areas were consistent with regions that are known to respond to novel stimuli (Ghazizadeh et al., 2020; Rolls et al., 2005). Of particular interest is the 45/46v cluster, which lay primarily in area 45 but had its peak in area 46v, and overlapped with our p46v ROI (**Figure 4B**). In the Ungrouped task for the same Low Exposure > High Exposure comparison, there was significant activity in 9/46, TEpv, TEO, and the caudate (FDRc < 0.05, height p < 0.005 unc., extent = 91; **Table 6**, **Figure 5**), all known to be novelty-sensitive regions as well (Ghazizadeh et al., 2020; Yamamoto et al., 2012). Taken together, the whole brain contrasts for Low Exposure > High Exposure in Grouped and Ungrouped demonstrate that known novelty-sensitive regions are more responsive to the low exposure images across both timing contexts.

**Figure 4.**
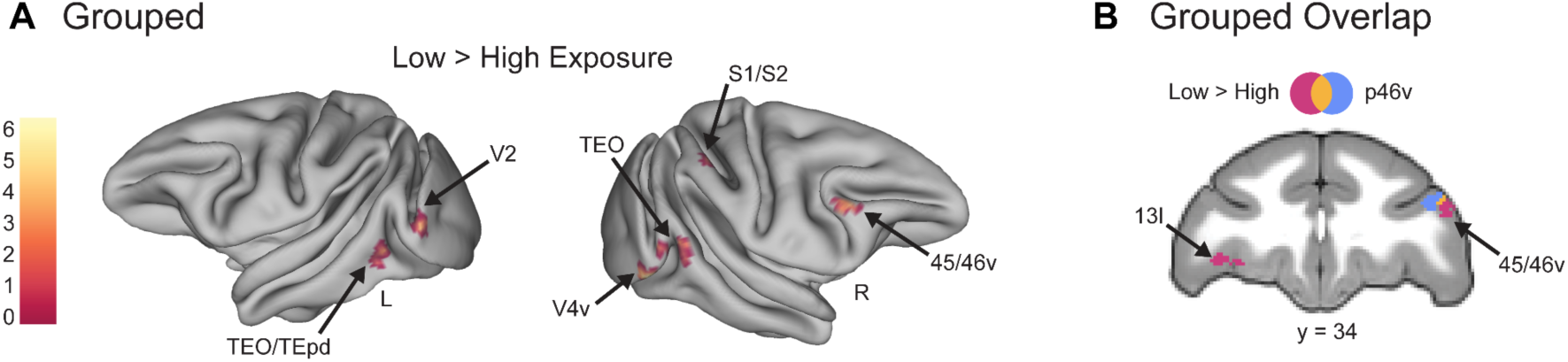
Whole brain contrasts show known novelty-responsive regions, including area IT, respond more to Low Exposure images in the Grouped task. Voxel-wise contrast of Low Exposure Images > High Exposure Images in the Grouped task false discovery rate (FDR) error cluster corrected for multiple comparisons (FDRc < 0.05, height p < 0.005 unc., extent = 116) is shown. **A.** Significant clusters in inferotemporal cortex (TEO/TEpd), visual area 2 (V2), ventral visual area 4 (V4v), somatosensory areas 1 and 2 (S1/S2), and ventral lateral prefrontal cortex (45/46v) are shown on inflated brains. Color bar indicates *t* value. **B.** Significant clusters in lateral orbitofrontal cortex (13l) and ventral lateral prefrontal cortex (45/46v) are shown in pink. The p46v ROI used in previous analyses is shown in blue and overlap between the contrast and ROI is shown in yellow.

**Figure 5.**
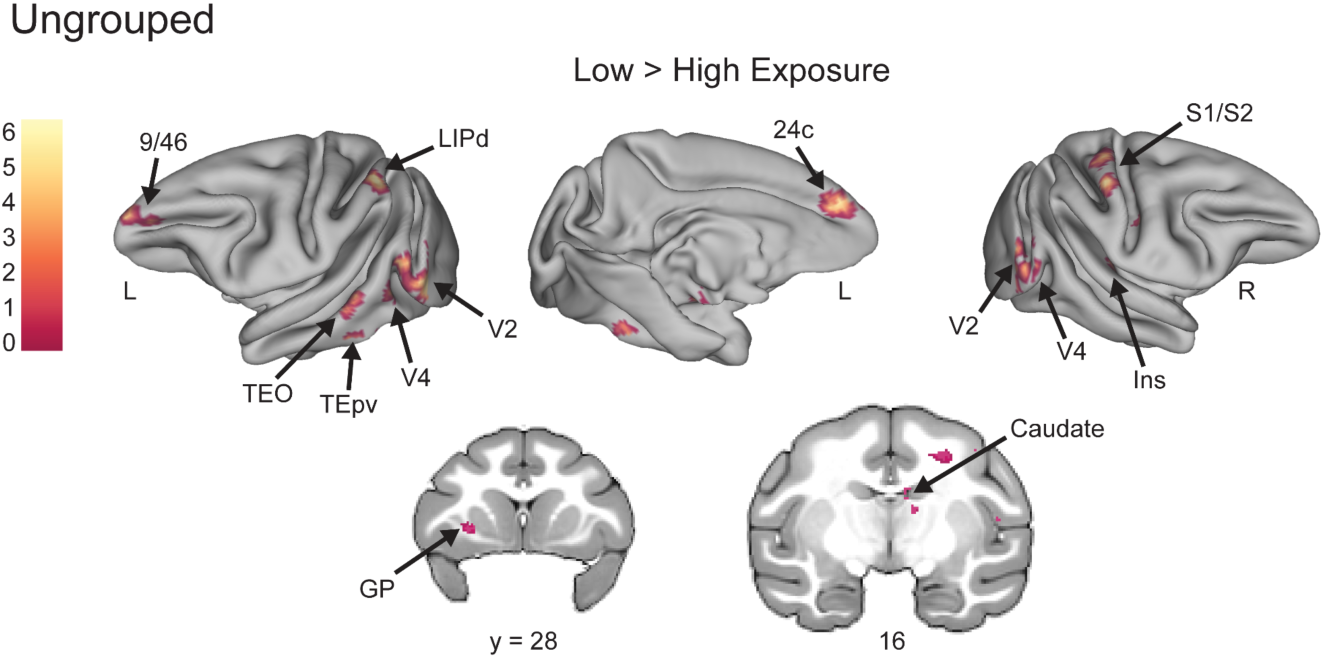
Novelty-sensitive temporal regions are active for Low Exposure stimuli in the Ungrouped task. Voxel-wise contrast of Low Exposure Images > High Exposure Images in Ungrouped task false discovery rate (FDR) error cluster corrected for multiple comparisons (FDRc < 0.05, height p < 0.005 unc., extent = 91) is shown. Significant clusters in prefrontal area 9/46, dorsal lateral intraparietal area (LIPd), inferotemporal cortex (TEO/TEpv), visual area 2 (V2), visual area 4 (V4), anterior midcingulate cortex (A24c), somatosensory areas 1 and 2 (S1/S2), Insula (Ins), globus pallidus (GP), and caudate are shown on inflated brains and coronal slices. Color bar indicates *t* value for inflated brain clusters.

**Table 6.** Coordinates of significant whole brain clusters from the Low Exposure > High Exposure contrast in the Grouped and Ungrouped tasks.

| **Contrast Location** | **Extent**<br>**(vox)** | **Peak**<br>**T-value** | **X** | **Y** | **Z** |
| --- | --- | --- | --- | --- | --- |
| **Grouped Low > High** |  |  |  |  |  |
| Lateral Area 13 | 116 | 5.14 | -13 | 34 | 14 |
| Area 45/46v | 156 | 11.33 | 19.5 | 33.5 | 23 |
| Areas 1 and 2 | 129 | 6.48 | 18.5 | 8.5 | 33.5 |
| Area TEO/TEpd | 309 | 7.65 | -28 | 3 | 6.5 |
| Area TEO | 263 | 7.29 | 30.5 | 1.5 | 17 |
| Ventral Visual Area 4 | 321 | 11.91 | 26 | -4 | 12.5 |
| Visual Area 2 | 171 | 6 | -29 | -5.5 | 17.5 |
| **Ungrouped Low > High** |  |  |  |  |  |
| Areas 9/46 & 24c | 293 | 6.89 | -4 | 40.5 | 21.5 |
| Globus Pallidus | 116 | 5.51 | -13 | 28 | 12.5 |
|  | 91 | 6.95 | 16 | 20 | 21.5 |
|  | 533 | 8.88 | 11.5 | 18 | 29.5 |
| Caudate Nucleus | 102 | 4.92 | 3 | 16 | 22.5 |
| Areas 1 and 2 | 133 | 7.39 | 19.5 | 13.5 | 32.5 |
| Granular Insula | 98 | 7.29 | 20.5 | 13.5 | 16.5 |
|  | 152 | 4.52 | -10 | 12 | 6.5 |
| Areas 1 and 2 | 157 | 8.23 | 14 | 9.5 | 36.5 |
| Area TPO | 173 | 5.56 | -24.5 | 8.5 | 13 |
| Dorsal Lateral<br>Intraparietal Area | 237 | 7.56 | -17 | 3 | 34 |
| Area TEpv | 195 | 12.76 | -22 | 2.5 | 4 |
| Area TEO | 278 | 6.48 | -28 | 1 | 14.5 |
| Dorsal Visual Area 4 | 355 | 5.96 | 25.5 | -1.5 | 24 |
| Visual Area 2 |  | 5.79 | 29.5 | -4 | 16.5 |
| Visual Area 2 | 98 | 7.62 | 25.5 | -7 | 14.5 |
| Visual Area 2 | 829 | 6.59 | -26 | -7 | 17 |
| Dorsal Visual Area 4 |  | 5.64 | -24.5 | -8 | 25.5 |

## Discussion

In this study, we investigated whether specific monkey area 46 subregions represent different levels of perceptual exposure to stimuli. We hypothesized that the sequence-monitoring subregion right p46f would not respond differently based on exposure, while adjacent subregion p46v would show distinct responses to different exposure levels. To test these hypotheses, we used an image-viewing task that presented High and Low Exposure images with both Grouped and Ungrouped timing. In support of our first hypothesis, area p46f did not show a significant difference in activation between levels of perceptual exposure. In support of our second hypothesis, the difference in responses to High and Low Exposure stimuli were greater in p46v than in p46f, showing a unique role of p46v in processing stimulus exposure. These results are consistent with the assertion that p46f responses may be specific to sequence deviants, as it does not display responses to Low Exposure stimuli that would indicate a role in broader novelty signaling. Additionally, they support a potential function of p46v in representing perceptual exposure of serially presented visual stimuli, emphasizing the remarkable functional specificity of adjacent area 46 subregions and serving to pinpoint exposure-dependent responses in LPFC.

We observed activation differences between subregions that indicate p46v may signal perceptual exposure, potentially mirroring canonical novelty responses, while p46f does not. Notably, we see an interaction between ROI and perceptual exposure level in the Grouped task, which appeared to be driven by increased p46v activation following Low Exposure stimuli. This modulation of activity in p46v supports a potential role for this region in signaling perceptual exposure, and the higher activation for Low Exposure stimuli mirrors the typical novelty response that appears in ventral LPFC (Ghazizadeh et al., 2020; Ghazizadeh & Hikosaka, 2022; Matsumoto et al., 2007). Additionally, this increase in activity to Low Exposure stimuli is not shared by p46f. Thus, our ROI analysis supports that perceptual exposure signals akin to novelty responses occur in p46v but not p46f, which is also indicated by our whole brain findings. When directly contrasting Low Exposure > High Exposure stimuli in the Grouped task, we found a significant cluster in area 45/46v, which partially overlapped with our *a priori* right p46v ROI and did not overlap with the p46f ROI. Together, these findings suggest that p46v, and not p46f, displays responses similar to canonical novelty effects when comparing Low and High Exposure stimuli.

While p46f did not display typical novelty responses, we did observe a marginal effect of perceptual exposure in p46f, such that the region showed numerically greater activation for High Exposure stimuli. Increased activation for more familiar stimuli has not been documented as frequently as novelty responses, although it has previously been recorded in monkey area IT (Anderson et al., 2008; Peissig et al., 2007) and human fusiform gyrus (Henson et al., 2000). In these studies, however, the stimuli used were highly familiar, and this long-term familiarity was proposed to underlie the effect (Peissig et al., 2007). Another explanation for the marginal effect we observe is that the High Exposure stimuli may have been associated with a sequential context. The High Exposure stimulus pool appeared as the habituation images for the sequential tasks not reported here. Their inclusion in the sequential task contributed significantly to the amount of exposure monkeys had with these stimuli, but also constitutes an additional context in which these images (and not the Low Exposure stimuli) were seen. While all responses analyzed here occurred during non-sequential tasks, it is possible that the High Exposure stimuli were associated to the sequential context and thus evoked sequence-related activity in p46f. Further study is required to determine whether p46f responds to stimuli with more perceptual exposure or if it may signal an association between the sequential context and individual images.

In addition to perceptual exposure differences, p46v also showed greater activation than p46f to visual stimuli overall. This result demonstrates another functional difference between these subregions, as p46v exhibits a greater sensitivity to visual stimuli regardless of exposure level. Area p46v has been proposed as a potential sensory integration hub based on the widespread functional connectivity patterns displayed by posterior 46, which shows correlation with a number of sensory regions (Rapan et al., 2023). The posterior bank of 46 also receives long range inputs from visual, visuomotor, and somatosensory regions (Barbas & Mesulam, 1985; Chavis & Pandya, 1976; Hackett et al., 1999). Most previous cortical tracing studies have not explicitly differentiated between the shoulder and fundus regions of posterior 46, however, and have instead largely focused on the cortical surface. Thus, it is unclear whether these sensory inputs project exclusively to p46v or to p46f as well. Our results illustrate response differences to sensory information between these regions, however, and emphasize the importance of distinguishing between the fundus and surrounding subregions.

In our whole brain analyses, we also found that activity evoked by Low Exposure stimuli corresponded to novelty-sensitive brain regions, suggesting that the degree of stimulus exposure was represented similarly to differences in familiarity. This similarity indicates that our stimulus pools were sufficiently distinct in their number of presentations to elicit familiarity or novelty-driven responses. Studies of familiarity and novelty often involve exposing animals to familiar stimuli over multiple weeks or months, while novel stimuli are presented either only once (entirely novel) or highly infrequently (Ghazizadeh et al., 2020; Peissig et al., 2007; Rolls et al., 2005; Xiang & Brown, 1998). We observed responses in novelty-responsive areas, including IT (Huang et al., 2018; Ranganath & Rainer, 2003; Zhang et al., 2022), ventral LPFC (Ghazizadeh et al., 2020; Ghazizadeh & Hikosaka, 2022; Matsumoto et al., 2007), orbital 13 (Ghazizadeh et al., 2020), and caudate (Yamamoto et al., 2012) while using session-unique stimuli that only differed in their degree of exposure five-fold, suggesting that these familiarity-dependent responses potentially arise earlier than has been previously observed in fMRI (Ghazizadeh et al., 2020). This observation is consistent with longitudinal electrophysiology results indicating that there are gradual changes in neural coding as stimuli shift from entirely novel to highly familiar as perceptual exposure increases, which has been demonstrated in LPFC (Rainer & Miller, 2000) and area IT (Koyano et al., 2023). As such, the High and Low Exposure stimulus pools can be conceptualized as existing at distinct points along a continuum of familiarity that mirrors the graded familiarity signal present in neural coding. The activation of novelty-sensitive regions by Low Exposure stimuli supports this concept, as the responses mirror those one would expect when using truly novel stimuli.

This study had the following limitations. First, it was limited in part by the no-report design of the task. While this design was chosen to avoid confounding the stimulus-related activity of interest with activity due to motor movements or decision making, it did limit the ability to determine whether the monkey had the capacity to use the information it gained about the stimuli being presented. Second, the current study used a dataset primarily designed to investigate sequence processing (Yusif Rodriguez et al., 2023). Though the analyses were designed to probe exposure level and did not include any data from sequential tasks, as previously discussed, it was possible that the presence of High Exposure stimuli in a sequential task could have influenced neural responses. While it is unlikely that a previous association with sequence is the primary driver of responses in a non-sequential task, we cannot rule out such effects. Therefore, dissociating stimulus exposure and context responses, particularly in known association areas such as area 46, is an important avenue of future research.

To summarize, we provided evidence that activity in a subregion of LPFC, p46v, signals differences in perceptual exposure of serially presented visual stimuli in a manner consistent with well-documented novelty responses across the brain. This perceptual exposure signal was not present in neighboring p46f, which is consistent with previous assertions that this subregion may perform a distinct and specific function in detecting sequence deviations. These findings advance our knowledge of the functional specificity of anatomically defined area 46 subregions and further localize perceptual exposure processing within LPFC.

## Data Availability

The data used in this study is available upon request. Correspondence should be addressed to.

## Author Contributions

Kyoko Leaman: Conceptualization, Formal Analysis, Writing - Original Draft, Writing - Review & Editing, Visualization. Nadira Yusif Rodriguez: Conceptualization, Methodology, Investigation, Writing - Original Draft, Writing - Review & Editing. Aarit Ahuja: Investigation. Debaleena Basu: Conceptualization, Investigation, Writing: Review & Editing. Theresa H. McKim: Conceptualization, Investigation. Theresa M. Desrochers: Conceptualization, Methodology, Investigation, Writing - Original Draft, Writing - Review & Editing, Supervision, Resources, Funding acquisition.

## Acknowledgements

This study was supported by the National Science Foundation (NSF) Established Program to Stimulate Competitive Research (EPSCoR) Neural Basis of Attention Grant 1632738 (N.Y.R. and T.M.D.), the National Institute of General Medical Sciences (NIGMS)-National Institutes of Health (NIH) Initiative to Maximize Student Development Grant IMSD R25GM083270 (N.Y.R.), the NIH-NIGMS Grant COBRE P20GM103645 (T.M.D), NIH National Institutes of Mental Health (NIMH) R21MH125010 (T.M.D.), NSF Faculty Early Career Development (CAREER) Program Award BCS-2143656 (T.M.D.), NIMH Research Project Grant R01MH131615 (T.M.D.), and the Carney Institute for Brain Science Innovation Award (T.M.D.). Part of this research was conducted using computational resources and services at the Center for Computation and Visualization, Brown University (NIH Grant S10OD025181).

We would like to thank Matthew Maestri for his help with animal training and data collection. We also appreciate the assistance and support of Dr. Michael Worden, Lynn Fanella, Fabienne McEleney, and Brown University’s MRI Facilities staff. We thank Dr. Lucija Jankovic Rapan, Dr. Nicola Palomero-Gallagher and Dr. Seán Froudist-Walsh for sharing the MEBRAINS atlas and regions of analysis used in this study. We also thank Dr. Katherine Conen and Hannah Doyle for their support throughout the development of this project, along with all members of the Sheinberg and Desrochers Labs that have contributed throughout this process.

